# DUX is a non-essential synchronizer of zygotic genome activation

**DOI:** 10.1101/569434

**Authors:** Alberto De Iaco, Sonia Verp, Sandra Offner, Didier Trono

**Affiliations:** School of Life Sciences, Ecole Polytechnique Fédérale de Lausanne (EPFL), 1015 Lausanne, Switzerland de Lausanne (EPFL), 1015 Lausanne, Switzerland

## Abstract

Some of the earliest transcripts produced in fertilized human and mouse oocytes code for DUX, a double homeodomain protein that promotes embryonic genome activation (EGA). Deleting *Dux* by genome editing at the 1- to 2-cell stage in the mouse impairs EGA and blastocyst maturation. Here, we demonstrate that mice carrying homozygous *Dux* deletions display markedly reduced expression of DUX target genes and defects in both pre- and post-implantation development, with notably a disruption of the pace of the first few cell divisions and significant rates of late embryonic mortality. However, some Dux^-/-^ embryos give raise to viable pups, indicating that DUX is important but not strictly essential for embryogenesis.

**Summary statement:** Murine DUX regulates transcription in the first embryonic cell divisions but it’s not necessary for embryogenesis

## Introduction

Fertilization of the vertebrate oocyte is followed by transcription of the parental genomes, a process known as zygotic or embryonic genome activation (ZGA or EGA) (Jukam et al., 2017). In zebrafish and Drosophila, maternally inherited transcription factors are responsible for this event (Lee et al., 2013; Liang et al., 2008), while in placental mammals the EGA transcriptional program is directly activated at or after the 2-cell (2C) stage by a family of transcription factors expressed after fertilization, the DUX proteins (De Iaco et al., 2017; Hendrickson et al., 2017; Whiddon et al., 2017). Recent studies suggest that DPPA2 and DPPA4 are maternal factors responsible in the mouse for DUX and downstream targets activation, although this model still needs to be validated *in vivo* (De Iaco et al., 2018; Eckersley-Maslin et al., 2019). Forced expression of DUX proteins in murine or human cell lines triggers the aberrant activation of EGA-restricted genes. Conversely, deleting *Dux* by CRISPR-mediated genome editing before the 2-cell stage in murine embryos leads to reduced expression of DUX targets such as *MERVL* and *Zscan4* and severe defects in early development, with many embryos failing to reach the morula/blastocyst stage (De Iaco et al., 2017). However, this procedure also yields some viable mice carrying heterozygous *Dux* deletions. Here, we demonstrate that crossing these Dux^+/-^ animals results in Dux^-/-^ embryos with impaired EGA and severe but not uniformly fatal defects in early development.

## Results and discussion

The murine *Dux* gene is found in tandem repeats of variable lengths in so-called macrosatellite repeats (Leidenroth et al., 2012). We injected zygotes collected from B6D2F1 mothers with sgRNAs directed at sequences flanking the *Dux* locus (Figure 1AB), and transferred the resulting products into pseudo-pregnant B6CBA mothers. One out of 42 pups carried a mono-allelic deletion of the targeted region (*Dux* ^+/-^). This animal was backcrossed twice with wild-type (WT) B6D2F1 mice to ensure germline transmission of the mutation. The resulting *Dux* ^+/-^ mice were healthy and did not display any macroscopic phenotype.

**Figure 1.**
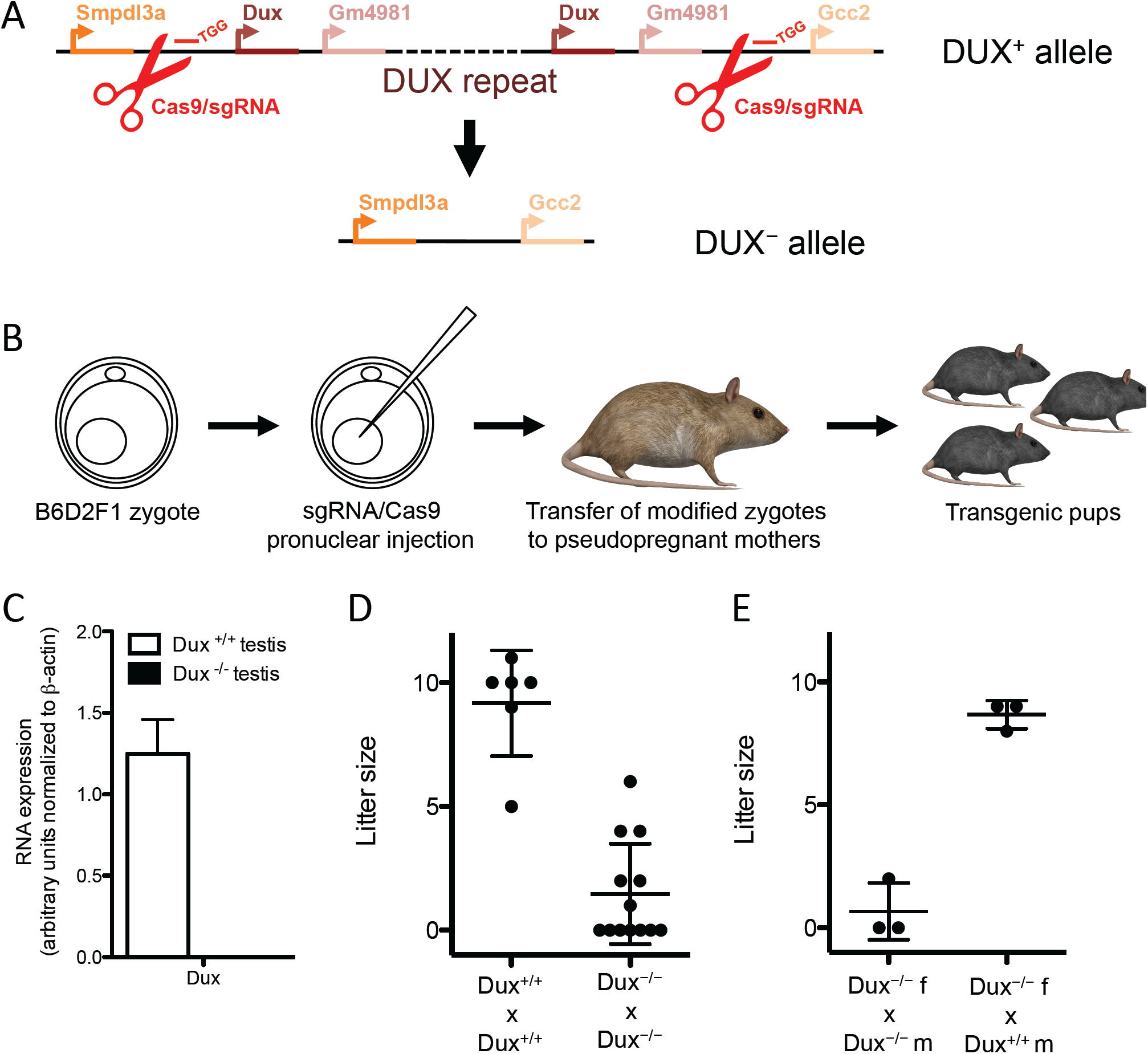
DUX promotes embryonic development but is not necessary for it. **(A)** Schematics of CRISPR/Cas9 depletion of *Dux* alleles. Single guide RNAs (sgRNA) targeting the flanking region of the *Dux* repeat recruit Cas9 nucleases for the excision of the allele. *Dux* and *Gm4981* are two isoforms of the *Dux* gene repeated in tandem in the *Dux* locus. Smpdl3a and Gcc2 are the genes flanking the *Dux* locus. **(B)** Generation of *Dux* ^-/-^ transgenic mice. Zygotes were injected in the pronucleus with plasmids encoding for Cas9 nuclease and the specific sgRNAs, transferred to a pseudopregnant mother and the transgenic pups were finally screened for the null alleles. **(C)** Expression of *Dux* in testes from adult *Dux* ^+/+^ and *Dux* ^-/-^ mice. **(D)** WT or *Dux* KO parents were crossed and litter size was quantified. **(E)** *Dux* ^-/-^ females were crossed with *Dux* ^-/-^ or *Dux* ^+/+^ males and litter size was quantified.

Transcription of *Dux* normally starts in zygotes just after fertilization and stops a few hours later (De Iaco et al., 2017), suggesting that the presence of a functional *Dux* allele is not necessary in germ cells. In our previous work, we demonstrated that inhibition of DUX expression in zygotes impairs early embryonic development. To characterize further the role of DUX, *Dux* ^+/-^ mice were crossed and the frequency of *Dux* mono- and bi-allelic deletions was determined in the progeny (Table 1). There was only a minor deviation from a Mendelian distribution of these genotypes, with a slightly lower than expected frequency of *Dux* ^-/-^ pups. Furthermore, adult *Dux* ^-/-^ mice were healthy and had a normal lifespan. To ensure that *Dux* was not expressed from some other genomic locus, the absence of its transcripts was verified in testis of *Dux* ^-/-^ mice, since this is the only adult tissue where these RNAs are normally detected (Snider et al., 2010) (Figure 1C).

**Table 1.** Genotype distribution from *Dux* ^+/-^ x *Dux* ^+/-^ crosses.

| Genotypes | $Dux^{+/+}$ | $Dux^{+/-}$ | $Dux^{-/-}$ |
| --- | --- | --- | --- |
| Observed n. of pups<br>(expected n. of pups) | 118<br>(109) | 225<br>(218) | 93<br>(109) |

To explore further the role of DUX in pre-implantation embryos, we compared the size of litters yielded by isogenic *Dux* ^+/+^ or *Dux* ^-/-^ crossings (Table 2, Figure 1D). Crosses between *Dux* ^-/-^ mice led to strong reductions in litter size and delayed delivery, and some of the rare pups were eaten by their mother after delivery, probably because they were either stillborn or exhibited physical impairments. Furthermore, some *Dux* ^-/-^ females failed to give any pup, even when crossed with *Dux*^-/-^ males that had previously demonstrated their fertility when bred with other *Dux*^-/-^ females (not illustrated). However, these apparently sterile *Dux*^-/-^ females produced litters of normal size following crosses with wild type males (Figure 1E).

**Table 2.** Genotype distribution from *Dux* ^+/+^ x *Dux* ^+/+^ and *Dux* ^-/-^ x *Dux* ^-/-^ Crosses.

| Crosses | $Dux^{+/+} \times Dux^{+/+}$ | $Dux^{-/-} \times Dux^{-/-}$ |
| --- | --- | --- |
| Total n. pups (litters) | 55 (6) | 36 (17) |
| Average litter size | 9.2 | 2.1 |
| Day of delivery (embryonic days) * | 19.5 | 20.8 |

We then analyzed whether the strong lethality observed after (*Dux*^-/-^ x *Dux*^-/-^) crosses occurred before or after implantation. For this, we repeated isogenic crosses of WT or *Dux* ^-/-^ mice, retrieved the zygotes at embryonic day 0.5 (E0.5, 27 embryos from 3 (WT x WT) and 42 embryos from 5 (*Dux*^-/-^ *x Dux*^-/-^) crosses), and monitored their *ex vivo* development for 4 days (Figure 2A). We found that up to E1.5 *Dux* ^-/-^ embryos divided faster that their WT counterparts yet sometimes unevenly, with formation of 3-cell (3C) structures. At E2.0, WT embryos caught up whereas *Dux* ^-/-^ embryos seemed partially blocked, to exhibit a clear delay at E3.5 with significantly reduced blastocyst formation. By E4.5, only 65% *Dux* ^-/-^ embryos reached the blastocyst stage, compared with 100% for WT. Confirming these findings, examination of E3.5 embryos from (WT x WT) or (*Dux*^-/-^ *x Dux*^-/-^) crosses revealed a strong delay in blastocyst formation and increased levels of lethality in the absence of DUX (Fig. 2BC). Finally, examining the uterus of *Dux* ^-/-^ females previously found to be sterile 18.5 days after crosses with *Dux* ^-/-^ males revealed a significant number of macroscopically normal embryos, suggesting that their apparent sterility was partly due to perinatal mortality (Figure 2D). In conclusion, a subset of embryos derived from *Dux* ^-/-^ crosses fail to implant, while the rest generally dies around birth.

**Figure 2.**
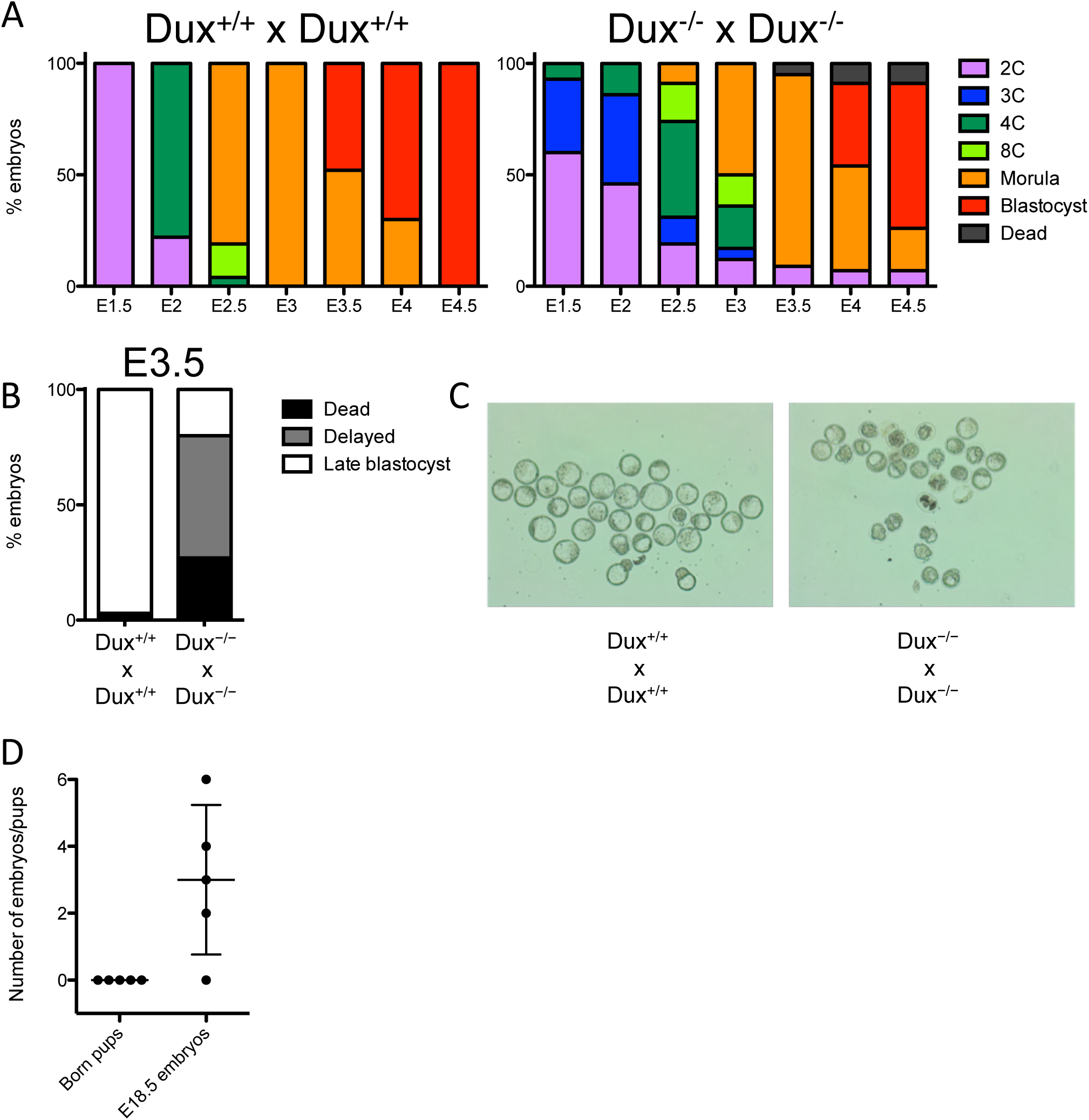
*Dux* promotes both pre- and post-implantation development. **(A)** Zygotes from *Dux* ^+/+^ (n = 3) or *Dux* ^-/-^ (n = 5) parents were monitored every 12 hours for their ability to differentiate *ex vivo* from embryonic day 1.5 (E1.5) to 4.5 (E4.5). Average percent of *Dux* ^+/+^ (n = 27) or *Dux* ^-/-^ (n = 42) embryos reaching a specific embryonic stage at each time point is represented. E3.5 embryos from WT (n = 30) or *Dux* KO (n = 28) parents were collected. **(B)** Average percent of embryos reaching the late blastocyst stages (white) or failing to differentiate (delayed embryos, grey; dead embryos, black) was quantified. **(C)** Bright-field images of the E3.5 embryos. **(D)** *Dux* ^-/-^ males and females were bred and number of born pups was quantified. The same animals were bred again and embryos were quantified at E18.5.

Finally, we tested the consequences of zygotic DUX on the transcriptional program of 2C-stage embryos. We collected 17 zygotes from three heterozygous *Dux* ^+/-^ x *Dux* ^+/-^ crosses, incubated them in vitro and collected RNA 5 hours after the formation of 2C embryos (Figure 3A). Three of these contained undetectable levels of *Dux* transcripts, indicating that they most likely were *Dux* ^-/-^, and an additional 3 displayed decreased levels of this RNA compared to the other 11. Interestingly, all 6 *Dux* RNA-depleted 2C embryos exhibited significant reductions in the expression of some (MERVL, *Zscan4, Eif1a, Usp17la, B020004J07Rik, Tdpoz4* and *Cml2*), but not all (*Duxbl, Sp110, Zfp352*) genes previously suggested to represent DUX targets (De Iaco et al., 2017). We then bred 2 *WT* and 3 *Dux* ^-/-^ females with males from the same genetic background, and compared transcription of putative DUX target genes in the resulting 2C embryos. Products of the *Dux* ^-/-^ x *Dux* ^-/-^ crosses displayed a clear decrease in the expression of a subset of candidate DUX targets (MERVL, Zscan4, *Eif1a, Usp17la, B020004J07Rik*), while others (*Tdpoz4, Cml2, Duxbl, Sp110, Zfp352*) were again unaffected.

**Figure 3.**
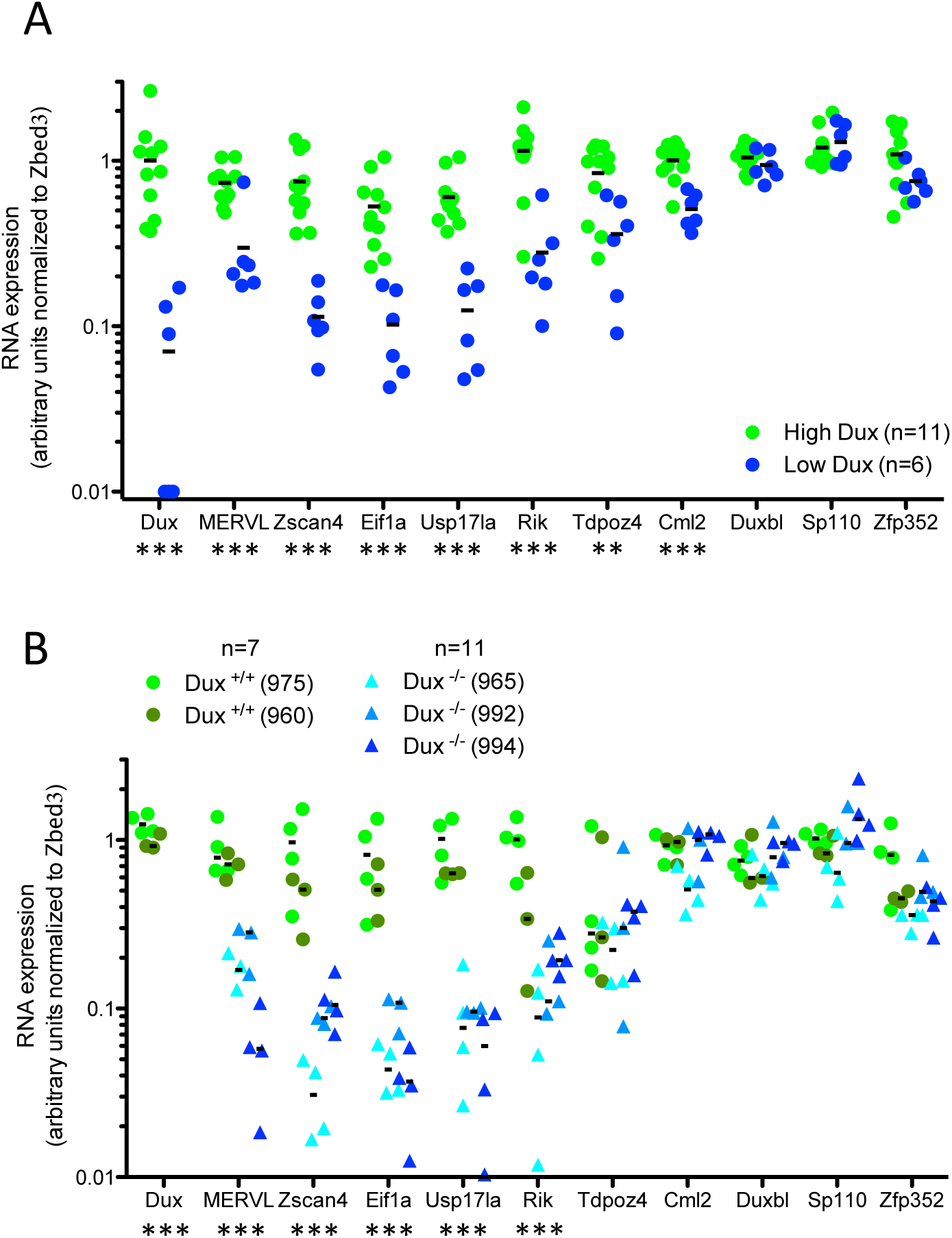
A subset of ZGA-specific genes is not expressed in 2C in absence of DUX. Comparative expression of *Dux*, early ZGA genes (*Zscan4, Eif1a, Usp17la, B020004J07Rik, Tdpoz4, Cml2, Duxbl, Sp110, Zfp352*), a 2C-restricted TE (MERVL), and *Zbed3*, a gene stably expressed during pre-implantation embryonic development, in 2C stage embryos derived from **(A)** *Dux* ^+/-^ breeding (n = 4) or **(B)** *Dux* ^+/+^ (n = 2) and *Dux* ^-/-^ (n = 3) breeding. Green and blue dots in (A) represent the mRNA levels of embryos expressing high or low levels of *Dux* respectively. Different shades of green or blue in (B) represent embryos collected from different mothers (975 and 960 are *Dux* ^+/+^ mothers, 965, 992 and 994 are *Dux* ^-/-^ mothers). Expression was normalized to *Zbed3*. ** p ≤ 0.01, *** p ≤ 0.001, t test.

In sum, the present work confirms that DUX promotes murine embryonic development. In spite of also surprisingly demonstrating that this factor is not absolutely essential for this process, it further reveals that DUX depletion results in a variable combination of pre- and post-implantation defects, the consequences of which additionally appear cumulative over generations. DUX-devoid embryos displayed deregulations in the timing and the ordinance of the first few cell divisions, various degrees of impairments in their ability to become blastocysts, and for those reaching that stage high levels of perinatal mortality. Nevertheless, these defects became truly apparent only at the second round of DUX-devoid embryogenesis, since the frequency of *Dux*^*-/-*^ pups derived from the crossing of heterozygous *Dux*^+/-^ parents was only slightly below a Mendelian distribution whereas the resulting *Dux*^*-/-*^ females yielded markedly reduced progenies, some even appearing sterile when crossed with *Dux*^*-/-*^ males. However, this defect was completely rescued by zygotic expression of *Dux*, since breeding these *Dux*^*-/-*^ females with WT males resulted in the production of normal size litters of pups devoid of obvious defects. Thus, the presence of DUX during only a few hours after fertilization appears to condition not only the conduct of the first few embryonic cell divisions, but also to bear consequences that extend well beyond the pre-implantation period, long after *Dux* transcripts have become undetectable. Deleting the *Dux* inducers *Dppa2* or *Dppa4* also results in perinatal lethality (Madan et al., 2009; Nakamura et al., 2011), but in this case defects in lung and skeletal development are observed, which correlate with the expression of these two genes later in embryogenesis. Future studies should therefore attempt to characterize better the molecular defects induced by DUX depletion, to explain how the full impact of the *Dux* KO phenotype is only expressed at the second generation, and how even at that point it can be fully rescued by paternally-encoded *Dux* zygotic expression.

## Material and Methods

### Plasmids

Two single guide RNAs (sgRNAs) targeting sequences flanking the *Dux* macrosatellite repeat (Figure 1A) were cloned into px330 using a standard protocol. The primers used to clone the sgRNAs are previously described (De Iaco et al., 2017).

### *Generation of transgenic mice carrying Dux* ^-/-^ alleles

Pronuclear injection was performed according to the standard protocol of the Transgenic Core Facility of EPFL. In summary, B6D2F1 mice were used as egg donors (6 weeks old). Mice were injected with PMSG (10 IU), and HCG (10 IU) 48 hours after. After mating females with B6D2F1 males, zygotes were collected and kept in KSOM medium pre-gassed in 5% CO2 at 37 °C. Embryos were then transferred to M2 medium and microinjected with 10 ng/μg of px330 plasmids encoding for Cas9 and the appropriate sgRNAs diluted in injection buffer (10mM Tris HCl pH7.5, 0.1mM EDTA pH8, 100mM NaCl). After microinjection, embryos were reimplanted in pseudopregnant B6CBA mothers. The pups delivered were genotyped for *Dux* null alleles using previously described primers (De Iaco et al., 2017). The mouse carrying the *Dux* null allele was then bred with B6D2F1 mice to ensure that the transgenic allele reached germ line and to dilute out any randomly integrated Cas9 transgene. This process was repeated once again to obtain second filial generation (F2) *Dux* ^-/+^ mice.

### Monitoring of pre-implantation embryos

Zygotes were collected and cultured in KSOM medium at 37 °C in 5% CO2 for 4 days. Each embryo was monitored every 12 hours to determine the stage of development.

Randomization and blind outcome assessment were not applied. All animal experiments were approved by the local veterinary office and carried out in accordance with the EU Directive (2010/63/ EU) for the care and use of laboratory animals.

### Standard PCR, RT-PCR and RNA sequencing

For genotyping the *Dux* null allele, genomic DNA was extracted with DNeasy Blood & Tissue Kits (QIAGEN) and the specific PCR products were amplified using PCR Master Mix 2X (Thermo Scientific) combined with the appropriate primers (design in Figure 1A) (De Iaco et al., 2017). Ambion Single Cell-to-CT kit (Thermo Fisher) was used for RNA extraction, cDNA conversion and mRNA pre-amplification of 2C stage embryos. Primers (previously listed) were used for SYBR green qPCR (Applied Biosystems) (De Iaco et al., 2017).

## Acknowledgements

We thank the Transgenic Core Facility of EPFL for technical assistance. This work was financed through grants from the Swiss National Science Foundation, the Gebert-Rüf Foundation, FP7 MC-ITN INGENIUM (290123), and the European Research Council (ERC 694658) to D.T.

## Author contributions

A.D.I and D.T. conceived the project and wrote the manuscript. A.D.I., S.V., S.O. designed the experiments, carried out the experiments and analyzed the data.

